# Efficient differentiation of vascular endothelial cells from dermal-derived mesenchymal stem cells induced by endothelial cell lines conditioned medium

**DOI:** 10.1101/271148

**Authors:** Ling Zhou, Xuping Niu, Jiannan Liang, Junqin Li, Jiao Li, Yueai Cheng, Yanfeng Meng, Qiang Wang, Xiaoli Yang, Gang Wang, Yu Shi, Erle Dang, Kaiming Zhang

## Abstract

**Objective:** To directionally-differentiate dermis-derived mesenchymal stem cells (DMSCs) into vascular endothelial cells (VECs) *in vitro*, providing an experimental basis for studies on the pathogenesis and treatment of vascular diseases.

**Methods:** After separation by adherent culture, VEC line supernatant, vascular endothelial growth factor (VEGF), bone morphogenetic protein-4 and hypoxia were used for the differentiation of VECs from DMSCs. The cell type was authenticated by flow cytometry, matrigel angiogenesis assay *in vitro*, and immunofluorescent staining during differentiation. The VEGF concentration was investigated by enzyme-linked immunosorbent assay.

**Results:** After 28 days of differentiation, the cell surface marker CD31 was significantly positive (80%-90%) by flow cytometry in the VEC line-conditioned culture, which was significantly higher than in the other groups. Differentiated DMSCs had the ability to ingest Dil-ac-LDL and vascularize in the conditioned culture, but not in the other groups. In the VEC line supernatant, the concentration of VEGF was very low. The VEGF concentration changed along with the differentiation into VECs in the medium of the conditioned culture group.

**Conclusion:** VEC line supernatant can induce the differentiation of DMSCs into VECs, possibly through the pathway except VEGF.

## Background

The abnormal function and/or quantity of blood vessels play a pivotal role in numerous diseases that significantly affect the patient’s quality of life. The high mortality associated with cardiovascular and cerebrovascular diseases is mainly caused by vascular disfunctions. Vascular disfunctions is also the vital characteristic of atherosclerotic peripheral artery disease, that affects more than 27 million people in North American and Europe [1]. Compared to drugs and traditional surgical treatments, cell therapy is an advantageous approach for the treatment of vascular abnormalities. To restore normal function, endothelial cells (ECs) are required in vascular tissue engineering or therapeutic angiogenesis [2]. However, an obstacle to the therapeutic use of ECs is the difficulty in obtaining enough cells (3]. Previous studies have suggested that endothelial progenitor cells (EPCs) may improve tissue perfusion in patients with myocardial infarction and peripheral arterial disease [4]. However, therapeutic uses of EPCs are limited due to their low proliferation capacity and the difficulty in obtaining a sufficient number of cells [3]. Therefore, it is necessary to search for new or alternative methods for the *in vitro* differentiation of stem cells into ECs, to enable their use in therapeutic angiogenesis.

Mesenchymal stem cells (MSCs) are adult stem cells derived from the mesoderm, with high self-renewal ability and multi-directional differentiation potential, abundant in organs and the interstitial connective tissue. It has been proven that this kind of cells can differentiate into ECs [1] and other types of mature cells, under certain conditions. Moreover, ECs derived from MSCs have better plasticity than adult ECs, as they can effectively upregulate markers associated with an arterial phenotype [5]. Multiple studies [6–8] have shown that MSCs are ideal seed cells for the *in vivo* and *in vitro* production of ECs for cell therapy.

The *in vitro* differentiation of MSCs into ECs is the primary step in cell therapy. To date, several approaches for generating ECs from MSCs have been established. Vascular endothelial growth factor (VEGF) is an important angiogenic signal protein. Oswald and colleagues reported that bone marrow-derived MSCs (BMSCs) were able to differentiate into endothelial-like cells in the presence of 2% fetal calf serum and 50 ng/ml VEGF [9]. Basic fibroblast growth factor (bFGF) can inhibit apoptosis of ECs under ischemic conditions by activating the endothelial cell hypoxia-induced factor-1 pathway [10]. It has been shown that more cells differentiate into ECs and participate in blood vessel formation in the presence of bFGF [1]. Data from Leroux *et al.* revealed that *ex vivo* hypoxic preconditioning of MSCs plays a unique role in the enhancement of vessel formation [11]. However, the efficiency of inducing MSCs to differentiate towards the endothelial cell lineage is limited [12].

In our previous work, we attempted to directionally-differentiate MSCs into ECs by hypoxia adding VEGF/stromal derived factor-1 (SDF-1) and gelatin. However, the differentiation never occurred, thus rendering these methods inadequate for use in cell therapy. In this study, we investigated the differentiation of dermis-derived MSCs (DMSCs) into ECs, by culture with vascular endothelial cell (VEC) line supernatant *in vitro*. It was found that in the conditioned culture, VECs were successfully differentiated, with high efficiency. Nevertheless, the differentiation by hypoxia, or in the presence of VEGF or bone morphogenetic protein-4 (BMP-4), were failed at the same time. This study presents a highly efficient, cost-effective method for the generation of VECs from DMSCs, suitable for differentiation on a large scale.

## Methods

### Subjects

In all, 10 Chinese Han volunteer controls (4 women and 6 men, mean age 34.2 years) were enrolled in this study. Volunteers exhibited no vascular diseases. Ethical approval for the experiments was obtained from the Medical Ethics Committee of Taiyuan City Centre Hospital, and all subjects provided informed consent.

### Isolation and expansion of DMSCs

The isolation and expansion of DMSCs were carried out as previously described [13]. The dermis was separated from the epidermis by incubating the samples in 0.25% dispase (Gibco, USA, lot NO.: 1563418) overnight at 4□. The remaining dermis samples were chopped further and collected in sterilized tubes containing DME/F-12(1:1) (Hyclone, USA, lot NO.: AC10248280) with 10% fetal bovine serum (Gibco, USA, lot NO.: 1908360C). The cell surface markers CD105, CD29, CD44, CD73, CD90, CD45, CD34 and human leucocyte antigen (HLA)-DR (Becton, Dickinson and Company, USA) were detected using a flow cytometer (Beckman Coulter Inc, USA), according to reference 13.

### In vitro differentiation of VECs from DMSCs

To differentiate DMSCs into VECs, the cells were divided into four groups (VEGF, BMP-4, hypoxic and conditioned culture group). The third passage of DMSCs involved the seeding at 1×10^4^ cells/cm^2^ on each well of a 24-well plate.

The VEGF group was cultured in basal medium supplemented with 50 ng/ml VEGF [7]. In the BMP-4 group, the cells were first differentiated to mesoderm using 20 ng/mL BMP-4, for 4 days, in human embryoid medium. Then, cells were attached to 0.67% gelatin-coated plates (1 well to 1 well ratio) and were cultured for an additional 10 days, in differentiation medium containing 50 ng/mL VEGF [14]. The hypoxic group was cultured in basal medium supplemented with 50 ng/ml VEGF for 14 days, in an incubator with a 2% O2 atmosphere [15]. EA.hy926 cells were obtained from the Shanghai Institutes for Biological Sciences, which was constructed by the fusion of the primary cultured human vein cells and the A549 clones of the anti-thioguanine under the duress of PEG. The fusant were grown on the HAT medium and screened by □ -factor correlation antigen. The VEC line supernatant was collected and stored at 4 □, then centrifuged at 300×g for 5 min [16] before use. The conditioned culture group was cultured with 50% VEC line supernatant and 50% fresh medium, containing 2% fetal bovine serum. The VEGF, BMP-4 and VEC line-conditioned groups were cultured in a humidified cell culture incubator at 37□, under 5% CO_2_, with close control of pH. The culture medium was changed every other day for all groups, and the cell morphology was also observed.

### Flow cytometry

To characterize the phenotype of VECs, flow cytometry was performed. Cells were collected at differentiation days of 0, 14, 21 and 28. After trypsinization and centrifugation, the cell samples were incubated with fluorescein isothiocyanate (FITC) -conjugated mouse anti-human CD31 (Beckman Coulter Inc., USA). Subsequently, the expression of CD31 on cell surface was analyzed by flow cytometry.

### In vitro matrigel angiogenesis assay

The potential of differentiated MSCs to form networks was examined using Matrigel Matrix Basement Membrane (Corning, USA, lot No.: 6172007), according to the manufacturer’s instructions.

The day before the experiment, the Matrigel was allowed to thaw at 4 □, overnight. With a refrigerated spearhead, 50μl Matrigel were added in each well of a 96-well plate. Then, the 96-well plates were incubated at 37 □, for 1 hour, to allow for solidification.

At day 0 and day 28, cells from four groups were trypsinized and plated at 1×10^4^ cells/cm^2^ onto Matrigel, and were then kept in a humidified cell culture incubator at 37 □, under 5% CO_2_, with close control of pH. Subsequently, the cells were examined under a light microscope with 20× objective magnification every 4 to 6 hours for 6 hours in all and the results were recorded.

### The uptake of acetylated- low-density lipoprotein (ac-LDL)

After differentiation, the medium was removed, and the cells were washed twice with PBS. Dil-acetylated low-density lipoprotein (Dil-ac-LDL) (Solarbio, China, Lot. No.: H7970) was incubated with the cells for 4 h, in a 37 □, 5% CO_2_ incubator. Cells were then washed twice with PBS and fixed with 4% paraformaldehyde for 5 min. After this, 2 mL of DAPI (Solarbio, China, Lot. No.20170412) were added in each well, for 10 min, at room temperature. After washing twice with PBS, fresh medium was added. The Dil-ac-LDL uptake of induced cells was observed by a fluorescence microscope.

### Secreted factors detection assay by enzyme-linked immunosorbent assay (ELISA)

Supernatant fluid from the VEC line-conditioned culture group was collected at differentiation days 0, 14, 21 and 28, and stored at −80 □. The ELISA (IBL, Germany, Cat.No.: F03060) was used to detect the level of VEGF in the supernatant. The VEC line supernatant was also collected for the detecting of VEGF, b-FGF, interleukin-5 (IL-5), IL-6, hepatocyte growth factor(HGF), transforming growth factor-β1(TGF-β1), SDF-1, granulocyte-macrophage colony stimulating factor(GM-CSF), Angiopoietin-1, soluble vascular cell adhesion molecule1(sVCAM-1), human insulin-like growth factor-1(IGF-1).

With zero blank, we measured the absorbance value (OD value) of each well, with 450 nm wavelength. According to the concentration of standard and corresponding OD value, the linear regression equation of standard curve was calculated. The concentration of different samples was calculated by the standard curve method. The final concentration is the actual measured concentration multiplied by the dilution factor.

## Results

### Cell morphology

In the initial days of culture, the cell morphology was long spindle, similar in all four groups (Fig. 1a–d). From culture day 18, the cells became gradually shorter in the VEC line-conditioned culture, whereas no changes were observed in the other three groups (Fig. 1e–h). Around day 28, the cells became short and stereoscopic in the conditioned culture, but remained long spindle-shaped in the other three groups (Fig. 1i–l).

**Fig. 1.**
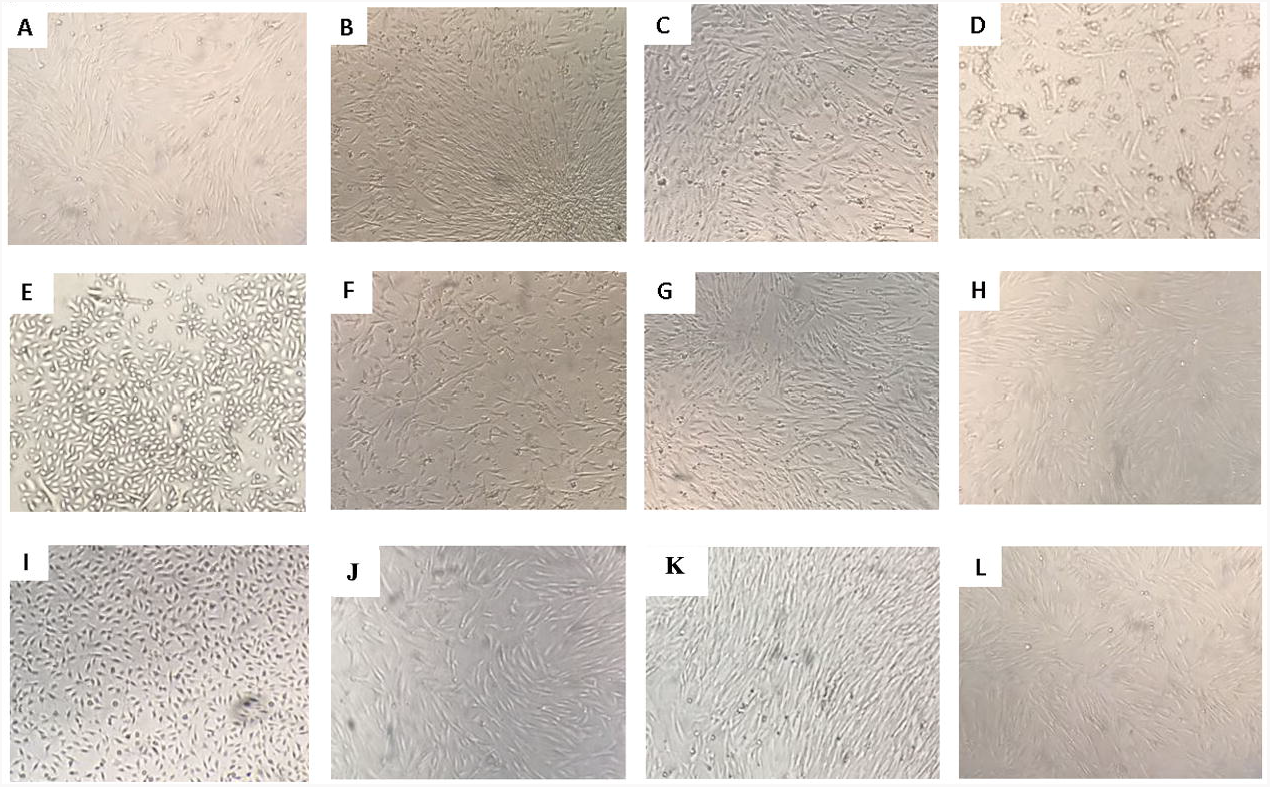
Cell morphology by inverted microscope (×100). **a** VEC line-conditioned culture group on day 0. **b** VEGF group on day 0. **c** BMP-4 group on day 0. **d** hypoxic group on day 0. **e** VEC line-conditioned culture group on day 18. **f** VEGF group on day 18. **g** BMP-4 group on day 18. **h** hypoxic group on day 18. **i** VEC line-conditioned culture group on day 28. **j** VEGF group on day 28. **k** BMP-4 group on day 28. **l** hypoxic group on day 28.

### Flow cytometry analysis

CD31 is a specific surface marker for VECs [1], thus the expression of CD31 was investigated by flow cytometer to detect the differentiation of VECs. In the initial days of culture, CD31 expression was extremely low in all four groups (Fig. 2a–d). At day 14, 21 and 28, the CD31 expression increased to 21.4%, 69.1% and 87.7%, respectively, in the VEC line-conditioned culture, but no change was detected in the other three groups (Fig. 2e–p, q).

**Fig. 2.**
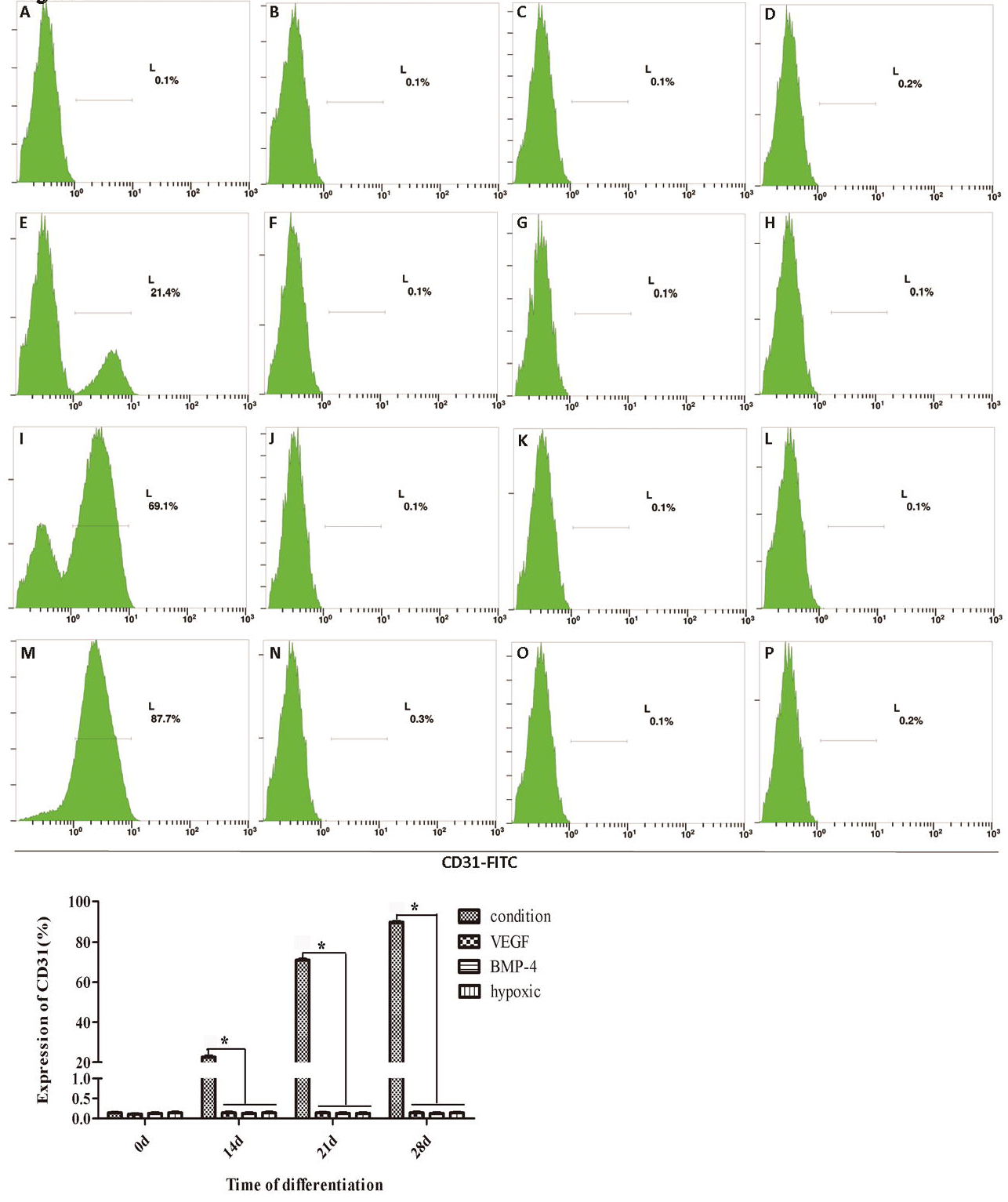
CD31 expression detected by flow cytometry. **a** VEC line-conditioned culture group on day 0. **b** VEGF group on day 0. **c** BMP-4 group on day 0. **d** hypoxic group on day 0. **e** VEC line-conditioned culture group on day 14. **f** VEGF group on day 14. **g** BMP-4 group on day 14. **h** hypoxic group on day 14. **i** VEC line-conditioned culture group on day 21. **j** VEGF group onday 21. **k** BMP-4 group on day 21. **l** hypoxic group on day 21. **m** VEC line-conditioned culture group on day 28. **n** VEGF group on day 28. **o** BMP-4 group on day 28. **p** hypoxic group on day 28. **q** CD31 expression in the four groups.

The difference examination was tested by SPSS, and the results show that there is a dramatic difference between VEC line supernatant group and other there groups at 14d, 21d, 28d (P<0.05). However, there is no difference among VEGF group, BMP-4 group and hypoxic group.

### Angiogenic network formation

In the initial days of differentiation, DMSCs were incapable of forming into tubes in the angiogenic network experiment, in either of the four groups (Fig. 3a). Around day 28, formed tubes were clearly visible in the VEC line-conditioned culture (Fig. 3b). However, no tubes were found in the other three groups (Fig. 3c-e).

**Fig. 3.**
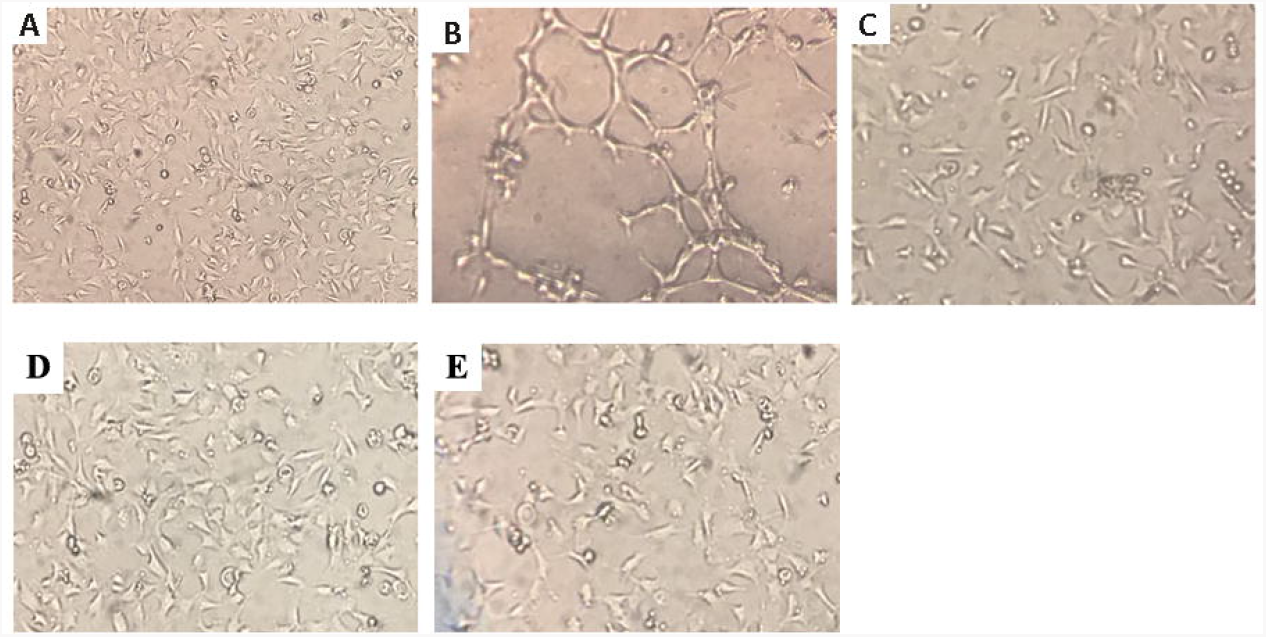
Angiogenic network formation (×100). **a** Four groups at day 0. **b** VEC line-conditioned culture at day 28. **c** VEGF group at day 28. **d** BMP-4 group at day 28. **e** hypoxic group at day 28.

### Dil-ac-LDL uptake

Only macrophages and ECs can uptake ac-LDL through scavenger receptors in the cell. Therefore, the ac-LDL uptake experiment is often used to identify VECs. The ac-LDL uptake ability was investigated in the VEC line-conditioned culture group after 28 days of differentiation. The bright field image is showed in the Fig. 4a. The red fluorescence is ac-LDL labeled by Dil (Fig. 4b), and the blue fluorescence is the nuclei labeled by DAPI (Fig. 4c). When the double fluorescence is stimulated at the same time, the cells display both red fluorescence and blue fluorescence, which proves that the ac-LDL labeled by Dil was uptook by the cells (Fig. 4d).

**Fig. 4.**
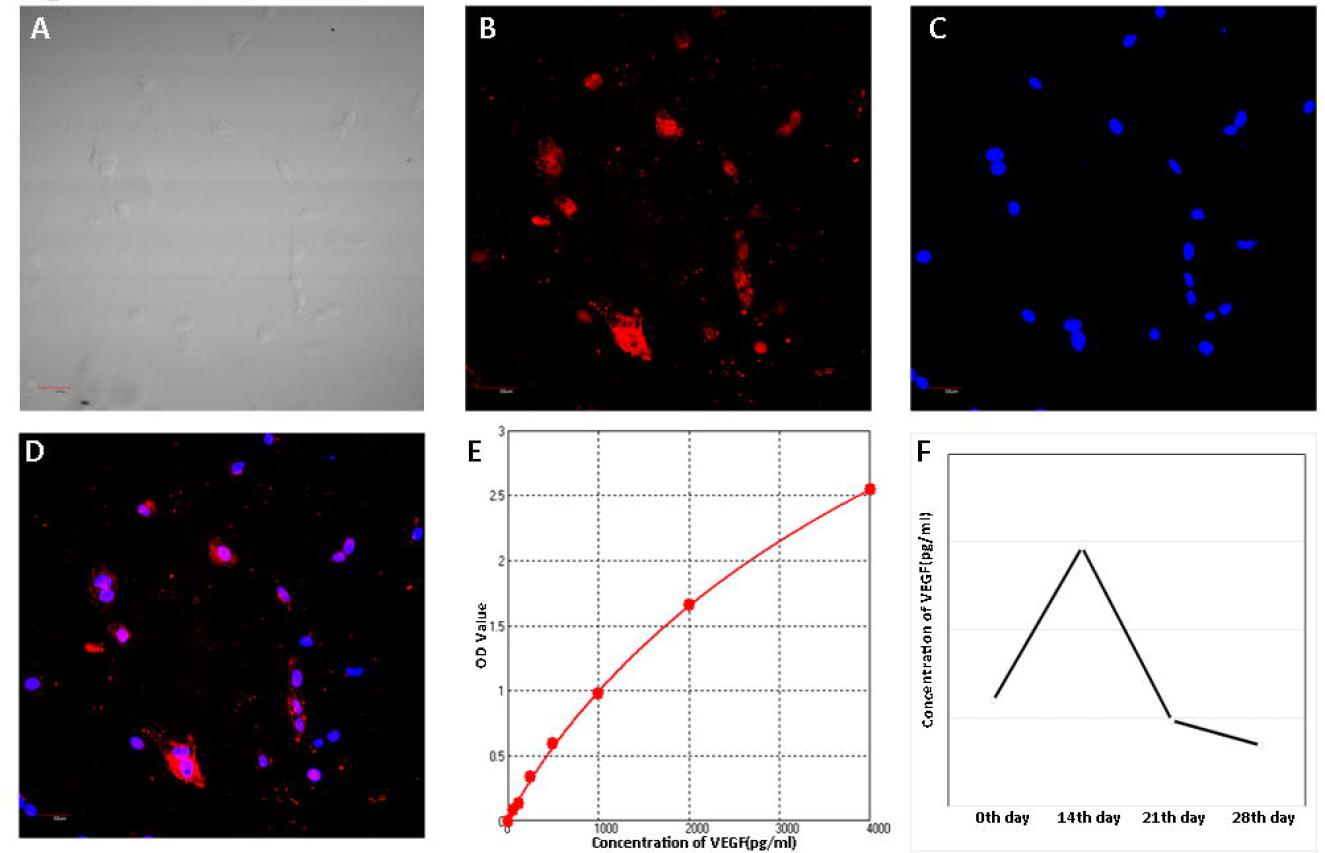
Dil-ac-LDL uptake and ELISA. **a** The bright field image (×200). **b** The uptake of ac-LDL labeled by Dil (×200). **c** Nuclei labeled by DAPI (×200). **d** The uptake of both ac-LDL and DAPI stimulated by double fluorescence (×200). **e** ELISA standard curve. **f** The concentration of VEGF at the different time points in the supernatant derived from the VEC line-conditioned culture group.

### VEGF detection by ELISA

The concentration of VEGF was investigated in the VEC line supernatant, the supernatant at differentiation days 0, 14, 21 and 28 in the conditioned culture, and the induction medium containing 50 ng/ml VEGF, by ELISA. The results showed that the concentration of VEGF in the VEC line supernatant was almost zero. The concentration of VEGF gradually increased in DMSCs during differentiation, and peaked to almost 3000 pg/mL, on the 14th day of differentiation. After reaching its peak, the concentration of VEGF started to decrease (Fig. 4e, f). The ELISA results showed that b-FGF(6.59 pg/ml), IL-5(34.62 pg/ml), HGF(33.26 pg/ml), TGF-β1(16.30 pg/ml), SDF-1(21.80 pg/ml), IGF-1(47.94 pg/ml) were rich in VEC line supernatant, especially for IL-6(62.28 pg/ml), GM-CSF(66.72 pg/ml), Angiopoietin-1(65.30 pg/ml), sVCAM-1(100 pg/ml).

## Discussion

The *in vitro* differentiation of ECs from MSCs is important for researching the pathogenesis of psoriasis, as well as for developing and assessing therapies for cardiovascular and cerebrovascular diseases. An increased number of blood vessels was observed in psoriatic lesions. Therefore, multiple methods of inducing MSCs to differentiate towards VECs have been used in our previous work, with the purpose of clarifying the relationship between DMSCs and angiogenesis in psoriatic lesion. Based on available knowledge, VEGF [7,17], SDF-1 [18], BMP-4 [14] and other factors were added into the differentiation medium, but the differentiation from DMSCs and BMSCs was not ideal in our research. In addition, Endothelial Cell Growth Medium-2 (EGM-2) (Lonza, USA, Lot.No.: 0000550876) [19,20], gelatin-coated culture plates [20] and low oxygen environment [1,15] were also used to induce differentiation. Unfortunately, the differentiation of VECs still failed. We sought to investigate the *in vitro* differentiation of DMSCs into VECs, for use in various research and therapeutic applications.

In this paper, we used a conditioned medium of VECs line (50% supernatant of VEC line and 50% fresh medium, containing 2% fetal bovine serum) to differentiate DMSCs into VECs. Compared to other differentiation approaches, the expression of CD31 in the VEC line-conditioned culture was significantly increased, which is a strong demonstration of VECs induced from MSCs. In the angiogenesis experiment, we found that the tubular structures form within 4-6 h after differentiation in the conditioned medium. However, no tubular structures were found in the VEGF, BMP-4 and hypoxic groups. After the Dil-ac-LDL uptake experiment, it was found that the cells had the ability to ingest Dil-ac-LDL after 28 days of induction in the conditioned culture, whereas no ac-LDL uptake was observed in the other groups. The abovementioned results proved that the induced cell type is VEC in the conditioned culture, but not in the other groups.

In this study, the more efficient *in vitro* VECs differentiation from MSCs was shown in the conditioned culture, compared to the other groups. However, the exact difference between the conditioned culture and the other groups attracted our attention. In most of the reported papers about VECs directional-differentiation *in vitro*, VEGF is a primary factor added to the differentiation medium, to ensure the successful generation of VECs, which regulate angioblasts from the mesoderm at the early developmental stage [21]. Therefore, we speculated that the concentration of VEGF in the supernatant of the VEC line was distinct from 50 ng/ml, which is more suited to the differentiation of VECs *in vitro*. However, the ELISA results showed that the VEGF content in the VEC line supernatant was very low, obviously different from 50 ng/ml. On the 14^th^ day of differentiation, when the differentiation of VECs was detected through CD31 expression on cell surface by FACS analysis, the VEGF concentration in the supernatant reached a peak. However, on the 28^th^ day of differentiation, the VEGF concentration was reduced to the same level as on day 0. The FACS results showed that most cells are MSCs on culture day 0, yet they differentiate into VECs by day 28. MSCs are known to secrete angiogenic growth factors like VEGF, bFGF and platelet derived growth factor (PDGF) for vascularization [22]. Our results agree with the VEGF secretion ability of MSCs. Moreover, the variant tendency of VEGF concentration prompted that the secretion ability of MSCs increase during their differentiation to VECs, and peak at the initial time of VEC differentiation.

This indicated that VEC-line supernatant with low level of VEGF could induce the VECs differentiation from MSCs *in vitro*. Therefore, we speculated that VEGF is not the only necessary factor for the differentiation of MSCs into VECs, which is in accord with the assumption of Ferguson *et al.* [23]. It must also be acknowledged that other growth factors were required for the differentiation process. The regulation of MSC activity by ECs has been investigated. Using EC-conditioned medium for MSC culture, Saleh and colleagues have shown that the paracrine signaling molecules secreted by ECs increase the proliferation and osteogenic differentiation of MSCs [24]. Another study demonstrated that co-cultured ECs secrete endothelin-1 (ET1) to upregulate the osteogenic and chondrogenic capacities of pre-differentiated MSCs, and the effects of ET1 on MSCs are mediated by AKT signaling [19]. ET1 is not the only soluble factor secreted by human aortic endothelial cells (HAECs) contributing to MSC regulation. Other molecules, such as PDGF, fibroblast growth factor, Wnt, bone morphogenetic protein, and Notch, have been reported to be involved in the regulation of MSC activities by ECs [24]. However, all of the reported regulation was related with ECs proliferation, osteogenic differentiation and adipogenic differentiation of MSCs. The effect of MSCs on VEC differentiation is yet not to be reported. VEC-line supernatant may induce VEC differentiation from MSCs *in vitro* through the same pathways of EC proliferation, osteogenic differentiation and adipogenic differentiation, as described above. Nevertheless, it is entirely possible other regulation mechanisms participate in the generation of VECs. Investigation into how the VEC-line supernatant regulates MSC differentiation through signaling pathways is beyond the scope of this study. This investigation shall be performed in future studies.

Moreover, the method only uses normal medium (DMEM), fetal bovine serum and supernatant of VEC-line, which were cheaper than other endothelial cell culture media and factors. The low cost of this inducing method could encourage its widespread application in the VEC differentiation *in vitro* for cell therapy.

## Conclusions

Overall, our study directionally-differentiate DMSCs into VECs, by culture with VEC line supernatant *in vitro*. Compared to the treatment with VEGF, BMP-4 or hypoxia, it was an efficient differentiation of VECs treated with VEC line-conditioned medium, which possibly through the pathway except VEGF. This study presents a highly efficient, cost-effective method for the generation of VECs from DMSCs, suitable for differentiation on a large scale.

## Abbreviations

DMSCs: Dermis-derived mesenchymal stem cells
VECs: Vascular endothelial cells
VEGF: Vascular endothelial growth factor
ECs: Endothelial cells
EPCs: Endothelial progenitor cells
MSCs: Mesenchymal stem cells
BMSCs: Bone marrow-derived MSCs
bFGF: Basic fibroblast growth factor
BMP-4: Bone morphogenetic protein-4
HLA: Human leucocyte antigen
FITC: Fluorescein isothiocyanate
Dil-ac-LDL: Dil-acetylated low-density lipoprotein
ELISA: Enzyme-linked immunosorbent assay
IL-5: Iinterleukin-5
HGF: Hepatocyte growth factor
TGF-β1: Transforming growth factor-β1
GM-CSF: Granulocyte-macrophage colony stimulating factor
sVCAM-1: Soluble vascular cell adhesion molecule1
IGF-1: Human insulin-like growth factor-1
EGM-2: Endothelial cell growth medium-2
PDGF: Platelet derived growth factor
ET1: Endothelin-1
HAECs: Human aortic endothelial cells.

## Ethics approval and consent to participate

Ethical approval for the experiments was obtained from the Medical Ethics Committee of Taiyuan City Centre Hospital, and all subjects provided informed consent.

## Consent for publication

Not applicable.

## Availability of data and material

All data generated or analysed during this study are included in this published article.

## Author’s contributions

LZ, JL and KZ designed the study and approved the final version. XN, YC collected the clinical data. XN, JL, QW, XY and YM performed the experiments and analyzed the data. LZ and XN assembled and interpreted the data, wrote the manuscript. JL and KZ are responsible for the integrity of this work and revised the manuscript. JL, QW, XY and ED assisted with cell culture, immunostaining and microscopy. GW, YS and KZ take responsibility for the integrity of the data and the accuracy of the data analysis. All authors read and approved the final manuscript.

## Acknowledgments

Not applicable.

